# Exploring the mechanisms governing the initiation of plexiform neurofibromas using *Prss56*^Cre^,*Nf1*-KO mouse model at a single-cell resolution

**DOI:** 10.1101/2025.07.04.662123

**Authors:** Myriam Mansour, Audrey Onfroy, Fanny Coulpier, Katarzyna J. Radomska, Pierre Wolkenstein, Piotr Topilko

## Abstract

About half of patients with the genetic disease Neurofibromatosis type 1 (NF1) develop benign nerve sheath tumors, called plexiform neurofibromas (pNFs). Despite important advances in understanding the pathogenesis of pNFs, mainly due to the plethora of dedicated mouse models, the mechanisms responsible for the initiation of this process remain poorly understood. Here, we used a *Nf1*-KO mouse model targeting biallelic loss of *Nf1* in boundary cap cells on a wild type and heterozygous *Nf1* background to explore the early events driving pNFs development. All mutants develop subcutaneous hyperplastic nerves with some progressing into pNFs from one year of age. We discovered that skin trauma accelerates this process, highlighting the role of inflammation. While skin trauma has no effect on control subcutaneous nerves, in the mutant nerves on a wildtype background, we observed an expansion of mutant Schwann cells expressing a profibrotic program. A similar response was observed in the mutant nerves on a heterozygous background without skin trauma, highlighting the impact of *Nf1* heterozygosity in the nerve micro-environment on the development of pNFs.

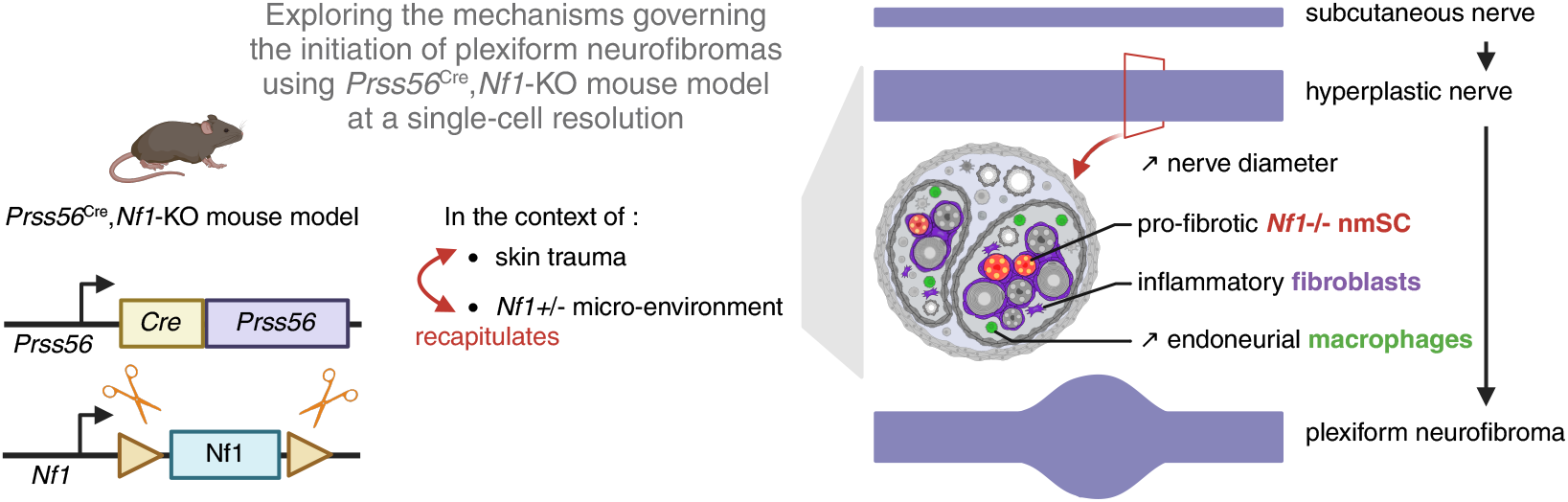

## Introduction

Neurofibromatosis type 1 (NF1) is a complex monogenetic disease due to mutations in the *NF1* gene encoding a negative regulator of the RAS signaling pathway (1). NF1 affects 1 in 3000 people worldwide and is characterized by a broad set of symptoms beginning in childhood and including café-au-lait macules, bone and eye lesions, learning disabilities, and neurofibromas (2). Neurofibromas (NFs) are benign nerve sheath tumors developing at the level of nerve roots (plexiform neurofibromas; pNFs) and nerve terminals of skin (cutaneous neurofibromas; cNFs). Both types of tumors originate from biallelic loss of *NF1* in Schwann cells (SCs) lineage (3). In addition to mutant SCs, NFs are composed of fibroblasts, a complex immune landscape, and are densely vascularized and innervated (4–6). Despite their common glial origin and benign character, pNFs and cNFs show differences. PNFs are developing prenatally or in early childhood in about 50% of NF1 patients while cNFs emerge in almost all patients starting from puberty (7). PNFs growth is slow but continuous and their number per patient is low while the number of cNFs may reach thousands and their growth is transient (2). Finally, about 10% of pNFs progress into malignant peripheral nerve sheath tumors (MPNSTs) while the malignant progression of cNFs was never reported (7, 8).

Mostly due to a plethora of *Nf1* genetically engineered mouse models, important progress was recently achieved in decoding cellular and molecular mechanisms driving the development of pNFs. In those models, biallelic inactivation of *Nf1* was targeted into the SCs lineage at successive stages of their development (9–12). Particularly, while loss of *Nf1* in SCs precursors, immature and non-myelinating SCs leads to the development of pNFs, inactivation of *Nf1* in myelinating SCs remains asymptomatic (13). Such an observation suggests that SCs in their immature state constitute the population at the origin of pNFs. Additionally, nerve trauma in mice with *Nf1* loss in myelinating SCs promotes their dedifferentiation and development of pNFs-like tumors further supporting the immature/inflammatory state of SCs as a prerequisite for pNFs development (14, 15). Recent studies showed that mutant SCs orchestrate the development of pNFs through their interaction with the complex nerve micro-environment, mostly composed of fibroblasts, immune cells, axons, and blood vessels (8, 16). One of the flagship molecules is stem cell factor (SCF) secreted by mutant SCs that binds to the receptor KIT on mast cells, which in turn stimulates collagen deposition through activation of fibroblasts by the transforming growth factor *β* (TGF*β*) (17, 18).

Despite such progress, the mechanisms governing the initiation of pNFs tumorigenesis remain to be decoded. In this study, we used the *Nf1*-KO mouse model, carrying biallelic *Nf1* loss and expression of fluorescent reporter Tomato targeted to a subpopulation of SCs originating from boundary cap (BC) cells, to address the early events and triggers responsible for pNFs development. We discovered that skin trauma and *Nf1* heterozygosity in the SCs micro-environment promote subcutaneous nerve enlargement defined as hyperplasia. While in the mutant nerves on the wild type (wt) background, the skin trauma is necessary to promote mutant SCs proliferation, *Nf1* heterozygosity in the nerve microenvironment in the absence of trauma is sufficient to mimic such an effect. Finally, we observed that mutant SCs express the pro-fibrotic marker periostin whose role is broadly known in the regulation of fibrosis and inflammation in different situations including cancer (19, 20).

## Results

### Skin trauma promotes subcutaneous nerve hyperplasia only in *Nf1* mutant mice

We have conceived a *Nf1* mutant mouse model (*Nf1*-KO) carrying simultaneous biallelic *Nf1* inactivation and permanent expression of fluorescent reporter Tomato (Tom) into *Prss56*-expressing BC cells and their derivatives, *Prss56* being a specific marker of BCs. BCs form clusters of stem-like cells transiently located during embryonic development at the dorsal root entry zone and ventral exit point of cranial and spinal nerves (21). Genetic fate mapping of BCs using the *Prss56*^Cre^ line identified derivatives migrating into the nerve roots and along spinal nerves into nerve endings of skin where they differentiate into myelinated and non-myelinated SCs (22). While in the nerve roots, almost all SCs originate from BCs, in the distal segments of spinal nerves (hypodermis, dermis, and epidermis) BC-derived SCs represent about 30% of the glia, the remaining cells originating from the neural crest (22). We have previously reported that *Nf1*-KO mice on wt; *Nf1*-KO^wt^ (*Prss56*^Cre^, *Nf1*^fl/fl^, *R26*^Tom^) and *Nf1* heterozygous; *Nf1*-KO^het^ (Prss56^Cre^, *Nf1*^fl/-^, R26^Tom^) background develop cNFs and pNFs starting from one year. PNFs are of two types, paraspinal, emerging at the level of dorsal root ganglia (DRG), and subcutaneous, developing along nerves in the hypodermis (Figure 1A) (23). It’s well known that the grouped house males, but not females, are naturally aggressive and have numerous skin bites. In the grouped house mutant males with skin trauma, we observed accelerated development of cNFs and subcutaneous, but not of paraspinal, pNFs (2).

**Fig. 1.**
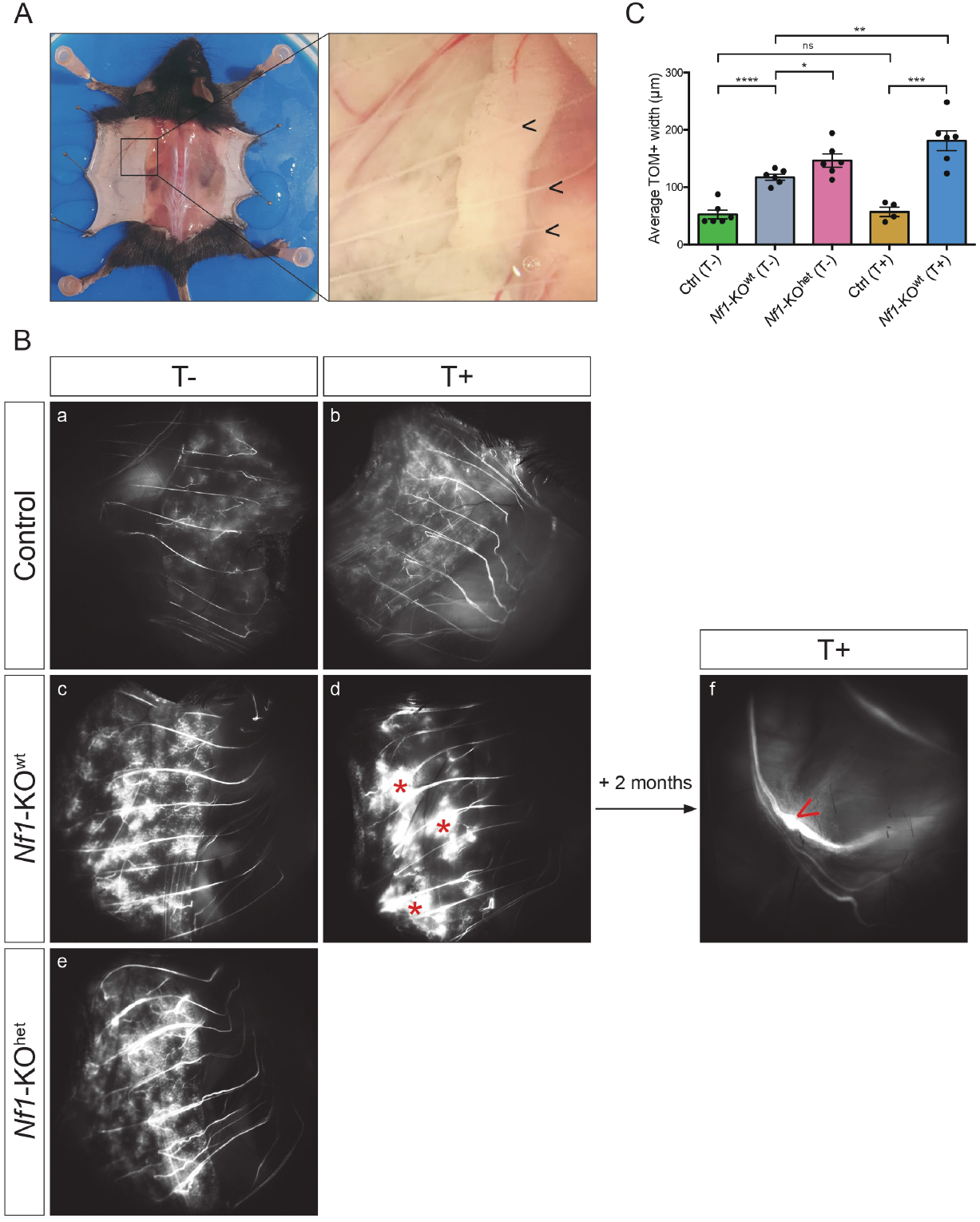
Macroscopic visualization of subcutaneous nerves from *Nf1*-KO mice. **A**. Metameric organization of nerves (marked by <). **B**. Macroscopic epifluorescent (TOM+) imaging of nerves from control, *Nf1*-KO^wt^ and *Nf1*-KO^het^ 3-month-old male mice, kept separately (without trauma; T-) or grouped house (with trauma; T+). Mutant nerves are enlarged with an increased proportion of TOM+ cells. Note the presence of numerous cutaneous neurofibromas (*) in grouped house mutant males. In the context of trauma, progression into pNF (marked by <) has been observed in 5-month-old *Nf1*-KO^wt^ mice. **C**. Average TOM+ nerve width, as mean by mouse +/-SEM (n = 6 nerves, 3-5 mice/experimental condition). Non-parametric t-test p-value: ns: non-significant, *: p-value < 0.05, **: p-value < 0.01, ***: p-value < 0.001, ****: p-value < 0.0001.

Since, in this model, *Nf1* inactivation takes place during embryogenesis and the first symptoms appear several months after birth, we characterized nerves from control and 3-monthold mutant males kept separately (without trauma; T-) or grouped house (with trauma; T+). Finally, we explored the role of *Nf1* heterozygosity in the nerve micro-environment by analyzing nerves from *Nf1*-KO^het^ males. All results presented in this study are based on analyses of at least 20 nerves per animal, distributed bilaterally and issued from 3 animals. First, observing nerves under the epifluorescent macroscopic showed no differences in the thickness of nerve segments populated by TOM+ cells across T- and T+ control mice, suggesting that skin trauma has no major impact on TOM+ SCs (Figures 1B-C). To validate such observations at the cellular level, we performed immunostaining of nerves from T- and T+ control mice with markers of all SCs (SOX10), all fibroblasts (PDGFR*α*), and subpopulations (SOX9, CLDN1, ZO-1), all immune cells (CD45) and macrophages (IBA1), the latter considered as a major immune component of injured nerves (Figure 2A) (5). No significant differences were observed in the cellular composition of control nerves supporting results from macroscopic analysis (Figure 2B).

**Fig. 2.**
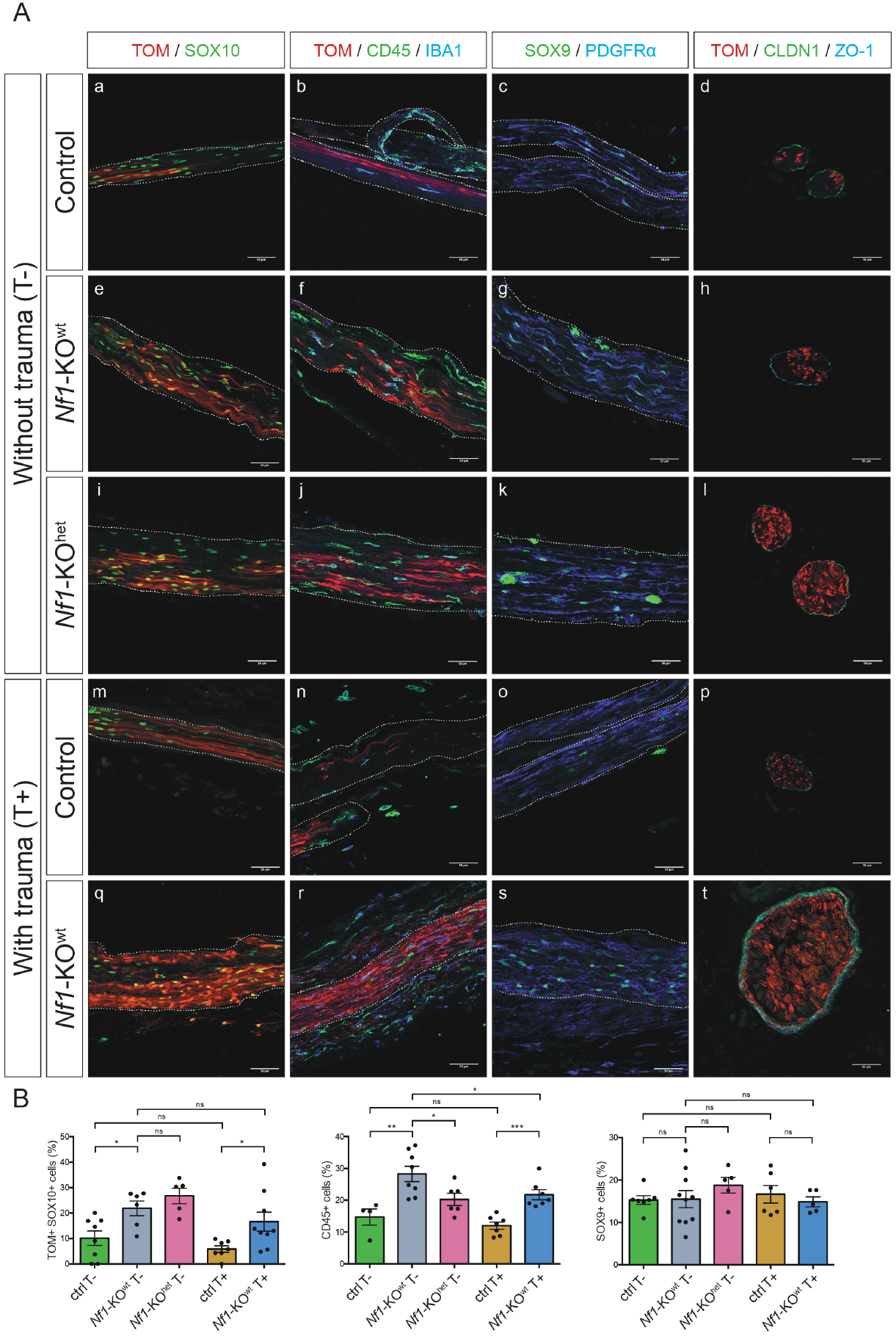
Immunohistochemical characterization of subcutaneous nerves. **A**. Immunofluorescence analysis of longitudinal and transverse sections of nerves from 3-month-old control, *Nf1*-KO^wt^ and *Nf1*-KO^het^ mice, kept separately (T-) or grouped house (T+), with markers of BC-derived Schwann cells (TOM), all Schwann cells (SOX10), all immune cells (CD45), macrophages (IBA1), all nerve fibroblasts (PDGFR^*α*^), endoneurial (SOX9) and perineurial (CLDN1 and ZO-1) cells. Scale bar: 50 *µ*m. **B**. Quantification of TOM+ SOX10+ Schwann cells, CD45+ immune cells, and SOX9+ fibroblasts in each experimental condition, over the total number of DAPI+ cells. Non-parametric t-test p-value: ns: non-significant, *: p-value < 0.05, **: p-value < 0.01, ***: p-value < 0.001, ****: p-value < 0.0001.

Then, we showed that the thickness of nerve segments populated by TOM+ cells is increased by 216% in T-*Nf1*-KO^wt^, by 334% in T+ *Nf1*-KO^wt^, and by 270% in T-*Nf1*-KO^het^ compared to control nerves (Figures 1B-C). We defined this local increase in nerve thickness as hyperplasia. Immunostaining of mutant and control nerves without trauma with a marker of proliferation (Ki67) did not identify any TOM+ dividing cells suggesting either rapid but transient proliferation of mutant SCs at the earlier period or low but constant proliferative activity, as previously reported (9). Immunostaining of mutant nerves (*Nf1*-KO^wt^ T-, *Nf1*-KO^wt^ T+ and *Nf1*-KO^het^ T-) with markers of SCs (TOM, SOX10) showed an increase of the TOM+ SCs proportion (Figures 1B-C). Such results suggest that while skin trauma in grouped house males has no effect on TOM+ SCs from control nerves, the same trigger promotes expansion of TOM+ mutant SCs in mutant nerves. Finally, in some *Nf1*-KO^wt^ males grouped house for more than 2-3 months, we observed the development of subcutaneous pNFs characterized by numerous tumor SCs detached from axons with abnormal multipolar morphology and decompacted perineurium, suggesting that hyperplastic nerves are at their origin (Figures 1B and S1).

Overall, our results suggest that in the *Nf1*-KO mice, skin trauma induced by bites and *Nf1* heterozygosity in the nerve micro-environment promotes local nerve hyperplasia with some progressing into pNFs.

### Skin trauma does not modify the transcriptomic signature of subcutaneous control nerves

To explore the changes in cell composition and transcriptomic activities that take place in *Nf1* mutant nerves, we performed single-cell RNA sequencing (scRNA-Seq) analysis of control and mutant nerves using 10X Genomics technology. We focused our analyses on the role of (i) *Nf1* loss in TOM+ SCs, (ii) *Nf1* heterozygosity in the nerve micro-environment, and (ii) skin trauma in grouped house males. To do so, nerves from control and *Nf1*-KO^wt^ and *Nf1*-KO^het^ males with and without skin trauma were collected, dissociated and isolated cells were processed for scRNA-Seq. Two replicates per experimental condition, except for T-*Nf1*-KO^wt^ (n=1), using 20 nerves/animal (3 animals/experimental condition, 60 nerves in total) were used and analyzed. Each biological replicate included between 6,000 and 14,000 cells. We added, to this set, two recently generated datasets from scRNA-Seq of subcutaneous pNFs issued from grouped house *Nf1*-KO^wt^ males (Radomska *et al*., In Press). All datasets were merged to form a transcriptomic atlas of nerves of 90,159 cells (Figure 3A). Since the cellular composition and transcriptomic signature for each cell type were recently established for the adult sciatic nerve (24, 25), we used this dataset as a reference to compare with our scRNA-Seq results from T- and T+ control nerves.

**Fig. 3.**
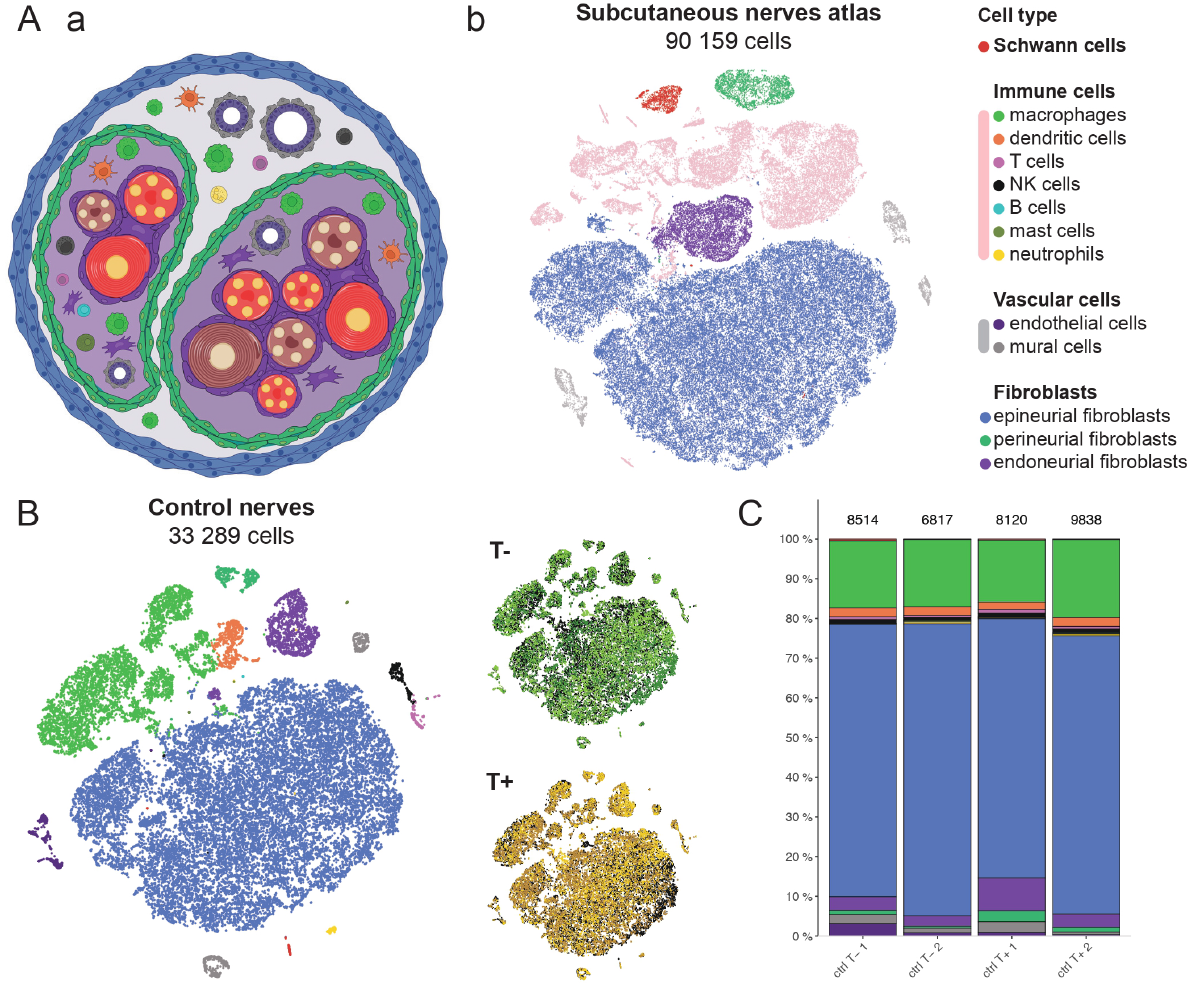
Transcriptomic atlas of control and *Nf1*-KO subcutaneous nerves. **A**. Single-cell nerve transcriptomic atlas (n = 11 independent datasets). **(a)** Schematic representation of nerve architecture including cell types. Created in BioRender. Coulpier, F. (2025) https://BioRender.com/hjczsp9. **(b)** tSNE visualization of 90,159 cells, colored according to cell type. Immune and vascular cells were colored in pink and gray respectively. See Figure S2A to visualize cell populations. **B**. tSNE presentation of 33,289 cells from control datasets (n = 4 independent datasets), colored according to cell types (left) or by experimental condition (right). In the right panel, each replicate (n = 2) within each experimental condition (T- or T+) is delineated as colored cells, while the background, corresponding to the other experimental condition, is represented as a black background. This tSNE is not corrected for sample effect. The right panel shows that all 4 samples (n = 2 per experimental condition) are perfectly overlapping, meaning there are no populations with unique transcriptomic signatures. **C**. Barplot showing cell populations representativeness in each control dataset. The text above bars corresponds to the number of cells in each dataset.

Both sciatic and control nerves include 13 distinct cell populations based on a cell specific markers based annotation (Figure S2D). The predominant population is made up of three types of nerve fibroblasts: epineurial (epiFb: *Clec3b*+, 70.8%), perineurial (periFb: *Tjp1*+, 0.8%), and endoneurial (endoFb: *Lum*^high^, 3.1%). The second major population corresponds to immune cells (*Ptprc*+) and includes macrophages (16.9%), dendritic cells (2.2%), T lymphocytes (0.6%), natural killer (NK) cells (0.8%), mast cells (0.2%), B lymphocytes (0.1%) and neutrophils (0.3%) (Figure S2C). The remaining populations correspond to endothelial cells (*Egfl7*+, 2.1%), mural cells (*Des*+, 1.8%), and rare SCs (*Plp1*+, 0.3%).

A subpopulation of SCs expresses *Tom* and corresponds to BCs-derived glia (Figure S3C). Such a low proportion of SCs is likely due to the incomplete nerve dissociation since we observed numerous TOM+ cells attached to myelin debris that were eliminated during the cell isolation process. All 13 cell populations were identified in the sciatic nerve, indicating that the global cell composition of adult nerves and sciatic nerve is the same. Our results further demonstrate that the nerves are still enveloped by the epineurium since this highly vascularized and composed of the rich connective tissue mantle nerve layer is no longer present in the distal part of the nerves entering the dermis.

Next, we explored the impact of skin trauma in control mice on cellular composition and transcriptomic signature of nerves (Figures 3B-C). To do so, we extracted, from the global dataset, 33,289 cells corresponding to nerves isolated from T+ and T-control mice. Then, we analyzed the intra-experimental condition heterogeneity by comparing two replicates issued from the same experimental condition and finally, we compared results across experimental conditions. Barplot for individual datasets showed that intra-experimental condition heterogeneity is slightly higher versus interexperimental condition differences (Figure 3C) revealing no major differences in cell composition between nerves isolated from T+ and T-control males. Immunolabelling on T+ and T-nerves with specific markers of SCs (TOM, SOX10), all immune cells (CD45), macrophages (IBA1), fibroblasts (PDGFR*α*, SOX9), perineurial fibroblasts (CLDN1, ZO-1) did not identify differences (Figures 2A-B).

Next, we compared the transcriptomic signature of each cell population between T+ and T-control nerves. First, we performed principal component analysis (PCA) and project cells using tSNE (Figure 3B), without applying sample-effect correction on the PCA space. Thus, tSNE was used to identify any new cell population based on overlapping between each dataset. No condition-specific population was identified showing that transcriptomic signatures for each cell population across T+ and T-control nerves are similar (Figure 3B).

Overall, our observations reveal that skin trauma induced by bites is probably too superficial to promote nerve injury and inflammation, in control males.

### Pro-fibrotic signature is a hallmark of *Nf1*-/-Schwann cells from hyperplastic nerves

Since *Nf1* mutant SCs are at the origin of the hyperplastic nerve phenotype, our initial focus was on this cell population. We extracted 1,046 SCs from the atlas and projected them using UMAP (Figure 4A). The dataset was annotated using a specific panel of SCs markers of myelinating (mSCs) and non-myelinating (nmSCs) subpopulations (Figure S3B). The majority of SCs issued from the nerves correspond to non-myelinating glia, which is consistent with the architecture of nerves, primarily composed of thin, poorly, or unmyelinated sensory axons (Figures 4Aa,C). Transcriptomic analyses led us to the following observations. First, there were no major differences in cellular composition and transcriptomic signature between *Tom*+ and *Tom*-SCs from the control nerves, showing that despite their distinct, BCs and neural crest, origin, they share a common molecular signature (Figures 4A,C, and S3A,C). Second, in the control nerves, the proportion of *Tom*+ SCs and their transcriptomic signature remained unchanged in the presence of trauma, confirming that the skin bites do not affect wt *Tom*+ SCs (Figures 4B-C and S3A,C). Third, no differences in transcriptomic signature were observed between wt and *Nf1*+/-SCs, demonstrating that *Nf1* heterozygosity in SCs in the absence of trauma does not affect their transcriptomic activities (Figures 4A,C and S3A).

**Fig. 4.**
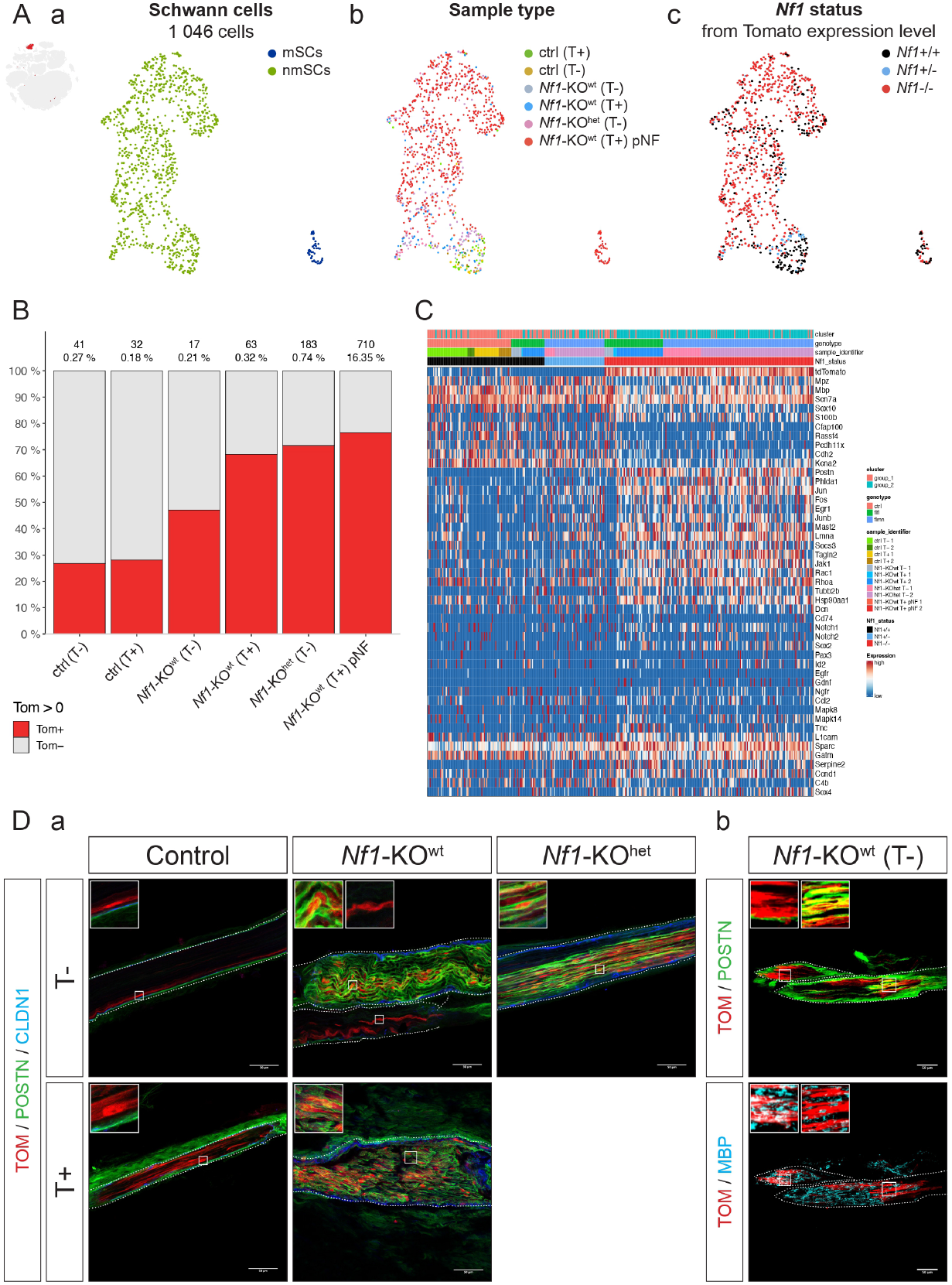
Transcriptomic signature of control and mutant Schwann cells from subcutaneous nerves. **A**. The iInset on the left represents all SCs, in red, extracted from the atlas. The 1,046 SCs are then represented on a UMAP colored by: **(a)** subtype, **(b)** experimental condition and **(c)** *Nf1* status, inferred from Tomato expression level (Figure S3C) and genotype. **B**. Barplot showing the ratio of BC-derived SCs (*Tom*+) versus neural crest derived SCs (*Tom*-) in each experimental condition. The number and proportion of SCs in the population among all cells within the experimental condition are indicated. **C**. Heatmap showing expression of genes in SCs, across experimental conditions. Gene expression is log-normalized, then scaled across cells. Cells from pNFs are not depicted in this heatmap. D. Immunostaining of longitudinal sections of control and mutant nerves with markers of **(a)** BC-derived SCs (TOM), periostin (POSTN), perineurial cells (CLDN1) and **(b)** myelin binding protein (MBP). Scale bar: 50 *µ*m. Abbreviation: mSCs, myelinating SCs; nmSCs, non-myelinating SCs.

Next, we compared *Nf1* mutant SCs isolated from T-nerves with those issued from T+ nerves and nerves on *Nf1*+/-genetic background. We observed that while *Tom*+ SCs represent 25% of all SCs in the control nerves (T+ and T-), this proportion is almost double (47%) in the T-mutant nerves and even higher (70%) in the presence of trauma or an *Nf1*+/-background (Figure 4B). This suggests that *Nf1* loss in SCs affects their proliferative activity, which is further enhanced by trauma and the *Nf1*+/-background. Bioinformatic analysis revealed that SCs from nerves are grouped in two sub-clusters (Figure S3D). Wt and *Nf1*+/-SCs grouped in subcluster 1 and were separated from subcluster 2 formed by mutant SCs issued from T+ and *Nf1*+/-background nerves supporting differences in their transcriptomic signature. Unfortunately, the small number of *Tom*+ mutant SCs from T-nerves on wt background (8 cells) does not allow us to classify them into one of the two subclusters. To further identify transcriptional differences between those 2 subclusters, we performed a differential expression (DE) analysis. Using two distinct DE methods, we identified 715 up-regulated and 27 down-regulated genes in subcluster 2 (Figure 4C). Such imbalance in the proportion of DE genes is likely due to globally higher transcriptional activity in *Nf1* mutant SCs. Among the top upregulated genes, we have identified a profibrotic factor periostin (*Postn*), *Phlda1*, and *Sox2* (Figures 4C and S3E). POSTN expression in control and mutant nerves was analyzed by IHC. In T- and T+ control nerves, POSTN expression is restricted to perineural cells (CLDN1+) and absent from TOM+ SCs. In mutant nerves, in addition to perineural cells, we observed a subpopulation of TOM+ SCs expressing high levels of POSTN (Figure 4Da). Finally, we showed that some mutant TOM+ SCs assembled in small clusters remain POSTN-. To differentiate between myelinating and non-myelinating SCs, we performed IHC with MBP, a marker for compact myelin. We observed that the TOM+, POSTN+ population corresponded to non-myelinating glial cells, while the TOM+, POSTN-expressed high levels of MBP and corresponded to myelinating SCs (Figure 4Db). These observations demonstrate that loss of NF1 in BC-derived SCs specifically activates expression of the pro-fibrotic marker POSTN in non-myelinating SCs without affecting myelinating SCs. A high level of periostin has also been observed in mutant SCs from pNFs, making this molecule a potential biomarker of the early development of pNFs (Figures S1d and S3E).

*Phlda1* encodes Pleckstrin Homology-Like Domain family A member 1 and emerges in many studies as a critical mediator of resistance to FGFR inhibition in cancer (26). Particularly, *Phlda1* and *Sox2* expressions are also maintained in mutant SCs from pNFs (Figure S3E). In addition to *Postn, Phlda1*, and *Sox2*, we observed an upregulation in mutant SCs from cluster 2 of several immediate early genes (IEGs) including *Fos, Jun, Junb*, and *Egr1* (Figure 4C). Since the expression of this set of genes was not observed in SCs from the control condition, we believed that such signature is rather associated with *Nf1* loss alone or in combination with trauma than with stress associated with tissue dissociation procedure (27). Moreover, overexpression of the same panel of IEGs in *Nf1* mutant SCs was also reported in pNFs from our model, further supporting their implication in the pathogenesis of those tumors. Finally, overexpression of *Postn, Phlda1, Sox2*, and the IEGs was also reported in SCs from the injured sciatic nerve and is associated with repairing SCs phenotype suggesting that *Nf1* loss in SCs in the presence of *Nf1*+/-background partly mimics such phenotype.

### Endoneurial fibroblasts from hyperplastic nerves over-activate the TNF signaling pathway

Peripheral nerves are protected by nerve fibroblasts. Based on their location in the nerve and functions, they can be classified into three distinct populations: epiFb, periFb, and endoFb. Since in pNFs, the nerve fibroblasts play a role in secreting the molecules of the extracellular matrix, we performed a comparative analysis of the proportions and transcriptomic activities of each type of fibroblasts between control and *Nf1* mutant nerves.

First, we focused on the epiFb population, which constitutes 87% of all nerve fibroblasts in our nerve atlas and forms a peripheral nerve mantle layer. 56,957 epiFb cells were extracted from the atlas (Figure 5Aa). The population of epiFb could be further divided into two subpopulations; A (orange) and B (green), distinguished by their specific gene expression profiles. The epiFb cells from subpopulation A express *Nav3, Calclr, Pla1a, Limch1*, and *Sema3e*, while those from subpopulation B are characterized by the expression of *Gdf10, Cxcl12* and *Cxcl14* (Figure 5Ba). Such two types of epiFb have not been described in the adult sciatic nerve (24). In the mutant versus control nerves comparison, we did not observe any differences in the proportion of epiFb and their transcriptomic signature, irrespective of the experimental conditions (Figure 5Bb). However, in pNFs from our mouse model, we mainly identified *Gdf10*+ epiFb belonging to the subpopulation B (Figure 5Bb). This suggests either enhanced proliferation of the B subpopulation, or A to B switch in transcriptomic signature of the subpopulation A.

**Fig. 5.**
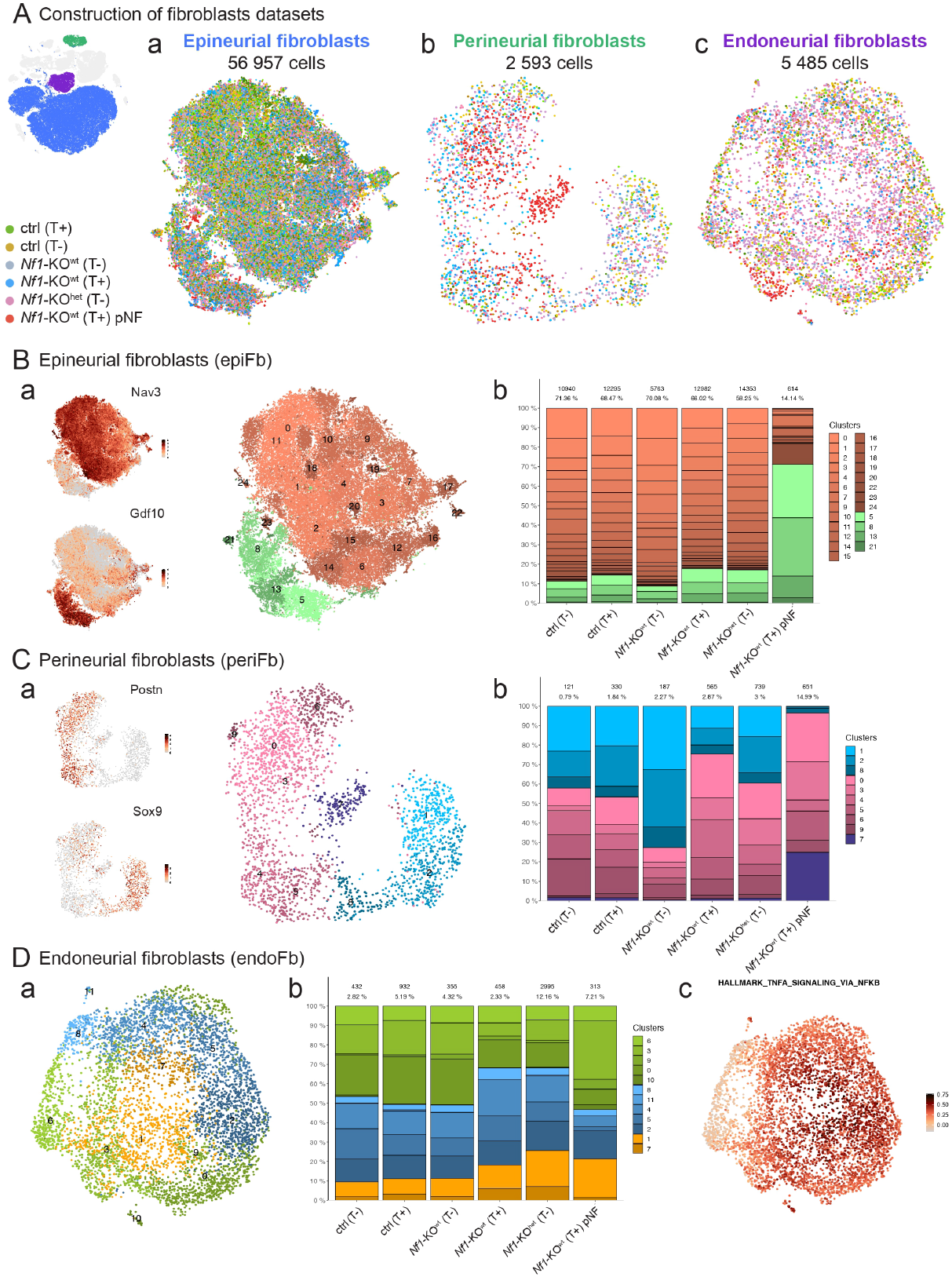
Transcriptomic signature of subcutaneous nerves derived fibroblasts. **A**. The inset on the left represents all fibroblasts from the atlas. 2D representation of each type of nerve fibroblasts including: **(a)** 56,957 epineurial fibroblasts (epiFb), **(b)** 2,593 perineurial fibroblasts (periFb) and **(c)** 5,485 endoneurial fibroblasts (endoFb) colored by experimental condition. **B**. Transcriptomic signature of epiFb **(a)** log-normalized gene expression levels identifying two subpopulations among epiFb. The orange population is *Nav3*+ while the green population is *Gdf10*+. **(b)** Barplot showing clusters representativeness in each experimental condition. The number and proportion they represent among all cells within the experimental condition are indicated. **C**. Transcriptomic signature of periFb **(a)** log-normalized gene expression level of two subpopulations of perineurial Fb. The blue population is *Sox9*+ while the pink population is *Postn*+. Purple subpopulation is specific to the pNFs dataset. **(b)** Barplot showing clusters representativeness in each experimental condition. The number and proportion they represent among all cells within the experimental condition are indicated. **D**. Transcriptomic signature of endoFb **(a)** endoFb clustering grouped into three subpopulations based on gene expression levels gradient. **(b)** Barplot showing clusters representativeness in each experimental condition. The number and proportion they represent among all cells within the experimental condition are indicated. **(c)** Representation of HALLMARK_TNFA_SIGNALING_VIA_NFKB genes set score, computed using Seurat’s AddModuleScore, showing score gradient from the surrounding cells to the middle yellow subpopulation.

The periFb population forms a protective barrier around nerve bundles, known as the perineurium. Its primary function is to protect nerves against pathogens and toxic compounds. We isolated and analyzed 2,593 periFb from our atlas, which accounted for 4% of the total fibroblasts (Figure 5Ab). In control nerves, periFb can be further divided into two subpopulations: A (blue) and B (pink). Subpopulation A is characterized by the expression of *Sox9*, while subpopulation B expresses *Postn* (Figure 5Ca). In the mutant nerves, we observed an increased proportion of both subpopulations of periFb compared to controls (Figure 5Cb). However, we did not observe differences in their transcriptomic signature, regardless of the experimental conditions suggesting that molecular characteristics of periFb are not affected in mutant nerves. IHC analysis of control and mutant nerves using ZO-1 and CLDN1, both markers of compact perineurium, revealed no differences further supporting transcriptomic data (Figure 2A). Furthermore, in the pNFs from our model only the population expressing *Postn* was present (Figure 5Cb). This observation suggests either a transcriptional switch from *Sox9*+ to the *Postn*+ population of periFb or an expansion of the *Postn*-expressing Fb.

The endoFb population infiltrates the nerve bundles and is the only population of fibroblasts in direct contact with SCs. To examine endoFb, we isolated and analyzed 5,485 endoFb from our atlas, accounting for 9% of all fibroblasts (Figure 5Ac). Notably, all endoFb are grouped in one cluster. However, we identified three distinct populations connected by partly overlapping transcriptomic signatures (Figure 5Da). Population A (blue) expresses higher levels of *Dbi, Gng5, Serf2*, and ribosome-protein coding genes while population B (green) shows higher expression of *Nsd2, Neat1, Rian, Cdk8*, and *Lars2*. Population C (yellow) over-expresses several IEGs in addition to *Tnfaip1-6* and *Nfkbia*, both hallmarks of the TNF signaling pathway (Figure 5Dc). While in T- and T+ control nerves, endoFb is composed of 45% of cells from population A, 45% of cells from population B, and 10% cells from population C, we observed expansion of the population C to 25% in T+ mutant nerves (Figure 5Db). Such expansion of endoFb with TNF signature was also observed in pNFs from our model.

### Subcutaneous nerves from control and mutant mice share the complex immune landscape

Immune cells represent the second most abundant cell type, after fibroblasts, of the nerves. To further analyze the transcriptomic signature of each immune cell type, we isolated and analyzed 19,922 immune cells from our atlas, corresponding to approximately 21% of all cells (Figure 6A). Each class of immune cell was identified based on the expression of a panel of specific markers (Figure 6B) and was present in the nerves across all experimental conditions (Figure 6C). Among the immune cells, the predominant group corresponded to M1 (*Plac8*+, *Chil3*+) and M2 (*Pf4*+, *Mrc1*+) macrophages. The M2 population comprised epineurial and endoneurial macrophages, each type depicted by a specific transcriptomic profile and location within the nerve (28) (Figure 6B). Epineurial macrophages express *Lyve1, Retnla* and *Clec10a* and are present in the epineurium. Endoneurial macrophages, expressing *Bcl2a1b, Dst, Creb5*, and *Dock4*, are in close contact with SCs and endoneurial fibroblasts. This latter population can be separated into M2 *Cx3cr1*+ and *Cx3cr1*-macrophages. While the overall proportion of macrophages remains constant under the different experimental conditions, we observed a notable increase in the proportion of M2 *Cx3cr1*+ macrophages in T+ mutant nerves and T-mutant nerves on *Nf1*+/-background (Figures 6C-D) but not in the mutant T-nerves on wt background. While *Cx3cr1*+ and *Cx3cr1*-macrophages co-exist within mutant nerves, in pNFs, all macrophages express *Cx3cr1*+ (Figure 6D). Despite the important complexity of the other immune cell types populating nerves, we did not identify major differences in proportion or transcriptome signature between control and mutant nerves.

**Fig. 6.**
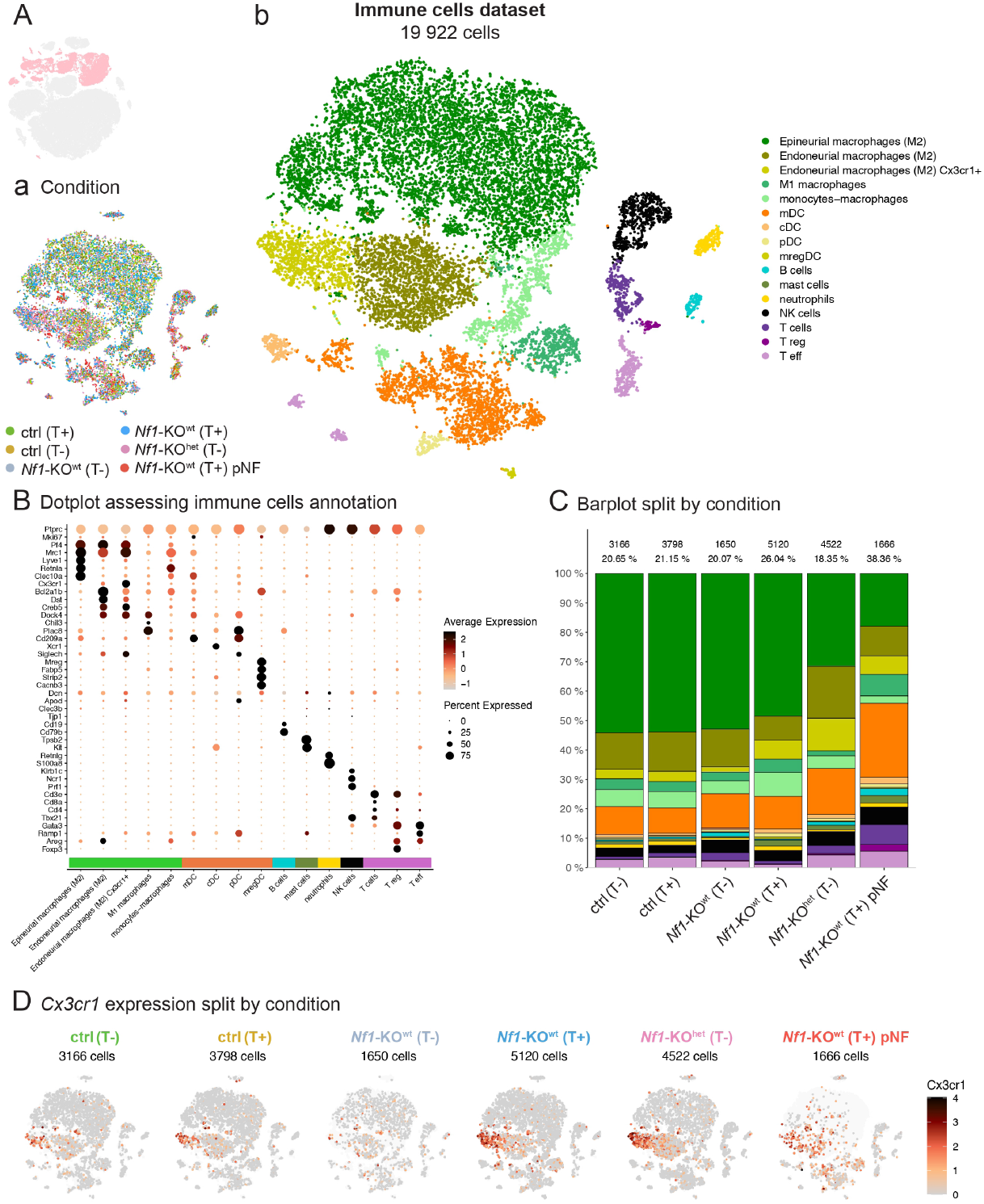
Figure 6. Transcriptomic signature of immune cells isolated from subcutaneous nerves. **A**. The inset on the left represents immune cells from the atlas. All 19,922 immune cells are represented on a tSNE colored by experimental condition **(a)** or by immune cell type **(b). B**. Dotplot assessing immune cell populations annotation. Dot size represents the proportion of cells expressing the corresponding marker. The dot color represents average gene expression levels for cells concerned by the dot. **C**. Barplot showing immune cell type representativeness in each experimental condition. The number and proportion they represent among all cells within the experimental condition are indicated. **D**. tSNE representations of *Cx3cr1* expression levels, for each experimental condition. The expression level is log normalized and ranges from gray to black. A light gray background represents cells from the other experimental conditions. Note that in pNFs datasets, the majority of macrophages and a subpopulation of dendritic cells expressed *Cx3cr1*.

## Discussion

Plexiform neurofibromas (pNFs) develop in about half of NF1 patients, but the mechanisms responsible for their initiation remain poorly characterized. In this study, we explored an *Nf1*-KO mouse model that exhibits *Nf1* bi-allelic loss restricted to BC cells and their derivatives. Adult *Nf1*-KO mice develop paraspinal and subcutaneous pNFs. We have shown that the development of subcutaneous pNFs is preceded by the presence of hyperplastic nerves characterized by the local accumulation of mutant SCs expressing high levels of periostin. Finally, we discovered that skin trauma in group-housed mutant males and *Nf1*+/-genetic background in the nerve micro-environment without trauma accelerated the development of pNFs but remained ineffective on paraspinal pNFs, highlighting the role of local triggers and *Nf1* heterozygosity in the nerve micro-environment as important players.

Our observations support the hyperplastic nerves origin of pNFs. First, hyperplastic nerves and pNFs share some commonalities, including an increased proportion of mutant SCs expressing Postn. Second, hyperplastic nerves are localized precisely where pNFs develop. Finally, the presence of numerous hyperplastic nerves but only a few pNFs suggests that their progression to pNFs is rare and probably requires additional local stimuli.

*Postn* expression is detected in mutant nerves and persists in pNFs, suggesting the role in their pathogenesis. Recently, *Postn* has been shown as one of the most upregulated extracellular matrix proteins in paraspinal pNFs, with a 7,38 fold change compared to wt DRGs (29). Known to contribute to immature SCs migration during embryogenesis (30), this protein is also expressed in repair SCs during nerve injury (31, 32). This highlights the immature or inflammatory character of mutant SCs in hyperplastic nerves and pNFs.

Single-cell level transcriptomic analysis of all SCs isolated from control nerves enabled us to compare for the first time the transcriptomic signature of *Tom*+ BC-derived versus *Tom*-neural crest-derived glial cells. We showed that both populations include mSCs and nmSCs, but no differences in transcriptomic signature were identified for these populations. This suggests that despite their distinct origins, BC-derived and NC-derived SCs share similar molecular characteristics. An analysis involving more cells is required to strengthen this observation.

Unexpectedly, we have shown that in control males, bite-induced skin trauma has no impact on the transcriptomic signature of nerves. No signs of mechanical nerve trauma, including nerve defasciculation and SC proliferation, were observed further supporting this observation. Contrastingly, under the same experimental conditions, mutant nerves behave differently since we observed an expansion in the proportion of *Tom*+ mutant SCs expressing *Postn* and a panel of IEGs. Since *Prss56*-expressing BCs give rise to nmSCs and mSCs in nerves, we wondered whether both types of mutant SCs responded to bite-induced skin trauma. We observed that despite their coexistence in the same nerve, only mutant nmSCs respond to skin trauma by proliferation and *Postn* expression.

Our observations suggest that mutant but not control nmSCs or mutant mSCs, respond to inflammation present in the injured dermis. Moreover, the fact that the transcriptomic signature of immune cells in mutant nerves upon skin trauma remains unchanged suggests that mutant nmSCs respond directly to such inflammatory stimuli.

In our model, hyperplastic nerves as well as paraspinal pNFs were often observed beneath cNFs, which develop in the dermis, reinforcing the role of local inflammation and suggesting that the mechanisms governing their initiation may be the same. Detailed comparison of the transcriptomic signatures of mutant SCs from pNFs and cNFs might help identify such mechanisms and associated candidate genes. By comparing the transcriptome of mutant nmSCs isolated from T+ and T-nerves, we hoped to identify genes whose expression is directly affected by *Nf1* loss. Unfortunately, the small number of SCs isolated from the T-mutant nerves made such an analysis impossible. However, periostin expression in T-mutant nerves but not in control nerves strongly suggest that expression of this profibrotic molecule is under NF1 control.

We observed that *Nf1* heterozygosity in the microenvironment of mutant nerves mimics the impact of trauma. This result was unexpected as we did not observe the increased incidence of pNFs in mutant males on *Nf1*+/-background. These results suggest that the *Nf1*+/-background might favor the initiation phase of nerve hyperplasia but is not sufficient for their progression into pNFs.

## Materials and Methods

### Animals

The following mouse lines were used and genotyped as described in the original publications (16, 22, 23). The mice with the following genotype were used: *Prss56*^Cre/+^, *R26*^tdTom/+^, *Nf1*^flox/flox^ (*Nf1*-KO^wt^), *Prss56*^Cre/+^, *R26*^tdTom/+^, *Nf1*^flox/-^ (*Nf1*-KO^het^) and *Prss56*^Cre/+^, *R26*^tdTom/+^ (control). The handling of the animals was conducted in agreement with French and European regulations. All experimentations on mice were approved by the French Ethical Committee (APAFIS #35296-2022020816327611). All mice were housed in a temperature- and humidity-controlled vivarium with a 12-hour dark-light cycle. Mice were genotyped by the PCR method.

### Sacrifice, dissection, and processing of subcutaneous nerves

Mice (males) were euthanized. Their skin was shaved, disinfected with 70% ethanol, and removed. The subcutaneous nerves were imaged under a Leica epifluorescence microscope. Pictures were taken using LaX software and images were analyzed using ImageJ software. Subcutaneous nerves were dissected and fixed by immersion in 4% paraformaldehyde (PFA) for 2 hours at 4°C on a shaking plate. After fixation, nerves were rinsed with 1X PBS before being cryoprotected by overnight immersion in 30% sucrose in PBS at 4°C. Nerves were placed in a Tissue-Tek OCT compound embedding medium (Sakura, ref: 4583) and were frozen at -20°C with dry ice and isopentane. The resulting blocks were stored at -80°C.

### Immunofluorescence

10 *µ*m frozen sections were blocked with 4% BSA in 0.3% TritonX-100 in 1X PBS for 1 hour, then stained with primary antibodies overnight at 4°C in the blocking solution. Secondary antibodies were added to the blocking solution for 2 hours at room temperature. Cell nuclei were labeled with Hoechst 33342 (Life technologies, ref: H3570). Sections were washed three times with 1X PBS between each step. Slides were mounted using Fluoromount G (Southern Biotec, ref: 0100-01). The following antibodies were used:

| Antibodies | Species | Type | Reference | Dilution |
| --- | --- | --- | --- | --- |
| tdTomato | Rat | Kerafast | EST203 | 1/1000 |
| mCherry | Goat | Sicgen | AB0040-200 | 1/1000 |
| RFP (DsRed) | Rabbit | Rockland | 600-401-379 | 1/1000 |
| SOX10 | Rabbit | Bio SB | BSB2584 | 1/500 |
| IBA1 | Rabbit | Wako | 019-19741 | 1/400 |
| PDGFRα | Goat | R&D systems | AF1062 | 1/100 |
| CD45 | Goat | BioTechne | AF114-SP | 1/100 |
| Ki67 | Rabbit | Abcam | ab15580 | 1/200 |
| ZO-1 | Rat | ThermoFisher | 14-9776-82 | 1/100 |
| CLDN1 | Rabbit | ThermoFisher | 71-7800 | 1/100 |
| PGP9.5 | Rabbit | Abcam | ab108986 | 1/200 |
| SOX9 | Rabbit | Abcam | ab185230 | 1/2000 |
| POSTN | Goat | BioTechne | AF2955 | 1/400 |
| TUJ1 | Chicken | Synaptic Systems | 302306 | 1/500 |
| CD49F | Rabbit | ThermoFisher | 710209 | 1/300 |
| S100B | Rabbit | DAKO | Z0311 | 1/500 |
| MBP | Rat | Millipore | MAB386 | 1/100 |

### Imaging

All images were acquired using the Leica TCS SP5 confocal microscope equipped with 10X, 25X, and 63X objectives. Images were saved in .tif format. All analyses (measurements, nerve delimitation, quantification) were done manually with ImageJ software. GraphPad Prism6 software was used for graphs. t-tests and nonparametric tests were used for statistical analyses.

### Single-cell RNA sequencing

#### Dissection and digestion

20 nerves per mouse were dissected and incubated in cold HBSS (Sigma, ref: H6648-500ML). After mechanical dissociation, nerves were transferred into a mix of enzymes (0.15% Trypsin (Sigma #T9201), 0.04% Hyaluronidase (Bioconcept #LS005474), 0.3% Collagenase type 2 (Bioconcept #LS004174) and DNAse I (25 µg/mL)) in HBSS, during 2 hours in a water bath. After digestion, nerves were tritured with a 1000 pipette, and a solution of 20% FBS in RPMI (Gibco, ref: 42401018) was added to stop digestion. Tubes were centrifuged and the supernatant was removed.

#### Cell sorting

Single viable cells were sorted (BD InfluxTM Cell Sorter) using a nuclear labeling DRAQ7TM Far-Red Fluorescent Live-Cell Impermeant DNA Dye (Abcam, ref: ab109202). Cells were collected in 20% FBS-RPMI solution.

#### Myelin depletion

Cells were centrifuged, the supernatant removed, and myelin depletion was performed by using anti-myelin beads (Miltenyi Biotec, ref: 130-096-733) in a 0.5% MACS solution (Miltenyi Biotec, ref: 5130-091-376). Cells were collected after the passage through the magnetic separator.

#### Single-cell RNA sequencing

Around 25,000 cells were loaded into one channel of the Chromium system using the V3.1 single-cell reagent kit (10X Genomics) to generate single-cell GEMs. Following capture and lysis, cDNAs were synthesized, and then amplified by PCR for 12 cycles as per the manufacturer’s protocol (10X Genomics). The amplified cDNAs were used to generate Illumina sequencing libraries that were each sequenced on one flow cell Nextseq500 Illumina.

#### Bioinformatic analyses

BCL files were processed using 10X Genomics Cell Ranger 7.0.1. Reads were mapped onto a custom mouse transcriptome based on GRCm39 release 108 transcriptome, in which we added sequences such as tdTomato. For both custom transcriptome construction and BCL to count matrix pipelines, we used Nextflow (33) and Singularity containerization (34). Count matrices were processed from raw matrices to the figures by applying the twelve tips for reproducible scRNA-Seq analysis (35). We used R Markdown notebooks to generate fully traceable analyses. A Singularity container containing R version 3.6.3 and all packages of interest was developed and used to compile notebooks. Reproducibility was ensured by rendering the notebooks at least twice and checking for the same outputs. The data, the code, the environment and the analysis results are available (See Code and Data Availability section).

Each dataset, corresponding to one biological replicate from one experimental condition, was analyzed individually from its raw count matrix using Seurat V3 package (36). Cells having less than 6 log number of UMI or less than 600 genes were filtered out. Doublet cells were removed using scDblFinder tool and scds in hybrid mode. Then, cells having more than 20% of UMI related to mitochondrial genes or more than 30% related to ribosomal genes were filtered out. The UMI count matrix for remaining cells was normalized using the LogNormalize method implemented in the Seurat V3 package. From the normalized count matrix, we used AddModuleScore to annotate cells for cell type using the following marker sets: *Sox10, Plp1, tdTomato, Nkain2, Ank3* and *Kcna1* for Schwann cells ; *Clec3b, Gdf10, Dpt, Gpx3, Slit3, Aebp1, Prrx1, Ly6c1, Pcolce2* and *Cxcl14* for epineurial fibroblasts (epiFb) ; *Lum, Smoc2, Spp1* and *Gpc3* for endoneurial fibroblasts (endoFb) ; *Cldn1, Tjp1, Itga6, Ildr2, Zbtb7c, Cttnbp2, Adamtsl3, Fxyd6* and *Itgb4* for perineurial fibroblasts (periFb) ; *Adgre1, Lyz2, Cd68, Pf4, Csf1r* and *Apoe* for macrophages ; *Cd209a, Mgl2, Tnip3, Mcemp1, Cfp, Xcr1* and *Napsa* for dendritic cells ; *Cd3g, Cd3d, Cd3e, Cd8a, Cd4, Foxp3, Gata3* and *Areg* for T lymphocytes ; *Nkg7, Klrk1, Klri2, Klrc2, Klrb1c, Ncr1* and *Klrb1f* for Natural Killer cells (NK cells) ; *Cd79a, Ly6d, Cd79b* and *Cd19b* for B lymphocytes ; *Tpsb2, Tpsab1, Cpa3* and *Kit* for mast cells ; *S100a8* and *S100a9* for neutrophils ; *Egfl7, Flt1, Ptprb, Shank3* and *Mecom* endothelial cells and *Des* and, *Rgs5, Notch3, Gucy1a1*, or, *Chodl, Nppc, Gal* and *Myf5* for mural cells. Note that two subpopulations of mural cells were identified, corresponding to the two sets of genes in brackets. Finally, cells were clustered using Seurat’s FindClusters function with a resolution of 2, on 20 principal components of a 100-dimension PCA obtained with Seurat’s RunPCA function from 3,000 highly variable features.

The same pipeline was used to either combine all 4 control datasets or all 11 datasets. Individual datasets were merged using base’s merge function. Genes expressed in less than 5 cells were removed. As for individual datasets, we generated a PCA. The sample-effect was removed on the PCA using the RunHarmony function from the harmony package (37). To generate UMAP or tSNE, we used the implementation for Seurat package: RunUMAP or RunTSNE. Cells were clustered on the harmonized PCA. To smoothen eventual misannotation, cell type annotation was grouped by cluster.

To build the 5 cell types-related datasets, the same procedure was applied. Cells were extracted from each individual datasets based on single cell or cluster smoothen annotation. After first processing as for the atlas, eventual contaminating cells were automatically removed through clustering and marker genes. A second processing was applied on the selected cells. For Schwann cells and fibroblasts datasets, sample-effect was removed from the normalized count matrix using the fastMNN function (38) from the bachelor package. For immune cells, we used harmony. Schwann cells projection was obtained using a diffusion map, implemented in the destiny package (39), on the corrected space, then UMAP. UMAP or tSNE from Seurat package were used for fibroblasts and immune cells datasets projection. Cells were clustered on the corrected space. Schwann cells were reannotated using the following genes: *Scn7a, Plac9, Ifitm3, Prkg1, Ptn* and *Ncam1* for nmSC and *Ncmap, Cldn19, Gfra1, Mlip, Fxyd6* and *Mpz* for mSC. Annotation was smoothed at a cluster level. Immune cells dataset was investigated for subpopulations based on clustering, differential expression and known markers from literature. For differential expression, both the methods implemented in DESeq2 package (40) and Wilcoxon test from Seurat’s FindMarkers function were used. Differentially expressed genes were filtered based on p-value adjusted, lower than 0.05.

## Graphical abstract

The graphical abstract has been Created in BioRender. Coulpier, F. (2025) https://BioRender.com/q3o0pis.

## Code and Data Availability

Single cell RNA sequencing data are available as FASTQ files and raw count matrices in ArrayExpress repository, under the accession number **E-MTAB-13334**. All the notebooks to process these data are available on GitHub. The GitHub Pages are activated to render the HTML pages (https://audrey-onfroy.github.io/Mansour_et_al_NF1_nerves_2025). The specific version of the analysis proposed here, corresponding to v1 on GitHub, is hosted on Zenodo under the record ID 15749911 (https://zenodo.org/records/15749911). The Singularity environment used to render the notebooks is available on Zenodo, under the record ID 15512051 (https://zenodo.org/records/15512051) for v1 and source files.

## Author Contributions

Conceptualization (AO, FC, PW, PT), Data Curation (AO), Formal Analysis (MM, AO), Funding Acquisition (PT), Investigation (MM, FC, KR), Project Administration (AO, FC, PT), Supervision (PW, PT), Validation (MM, AO, FC, PT), Visualization (MM, AO, FC, PT), Writing - Original Draft Preparation (MM, AO, FC, PT), Writing - Review & Editing (AO, PT)

## Supplementary Figures

Please refer to next pages for Figures S1, S2 and S3.

**Supplementary Figure 1.**
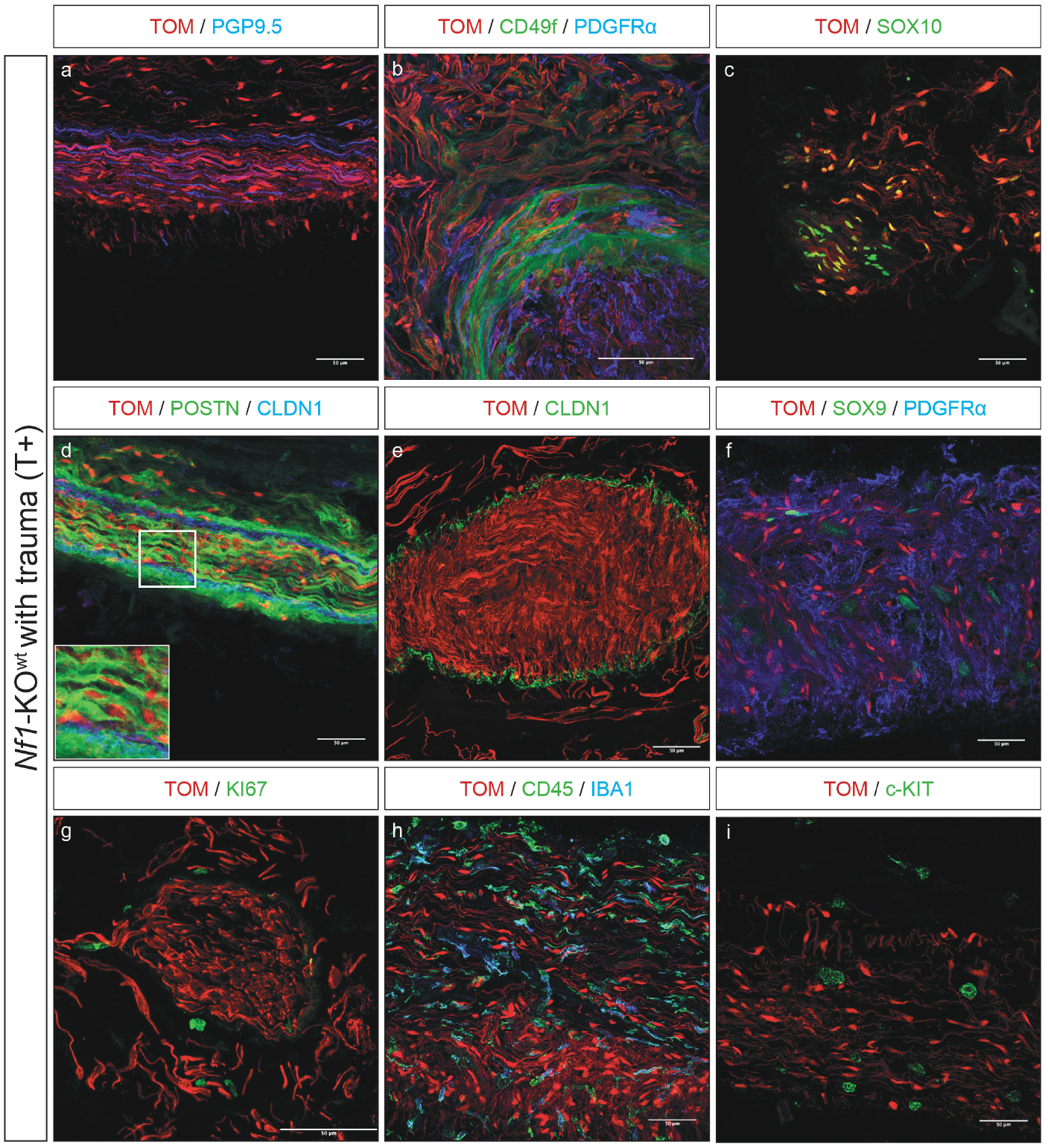
*Nf1*-KO^wt^ grouped house males develop subcutaneous plexiform neurofibromas. Immunofluorescence analyses of longitudinal and transverse sections of pNFs show accumulation of mutant SCs (TOM) that are detaching from axons (PGP9.5). When going out of nerve fascicles, SCs express CD49F (also known as ITGA6), which is also expressed by periFb. Once out, the CD49F expression shuts down. Mutant SCs still keep their glial identity (SOX10) and maintain expression of (POSTN), which is also found around perineurial cells (CLDN1). The perineurial layer is decompacted (CLDN1). pNF shows accumulation of fibroblasts (PDGFR*α*) including endoneurial (SOX9) fibroblasts. However, almost all cells are either quiescent or slowly proliferating (KI67). pNF are infiltrated by immune cells (CD45) including macrophages (IBA1). Scale bar: 50 *µ*m.

**Supplementary Figure 2.**
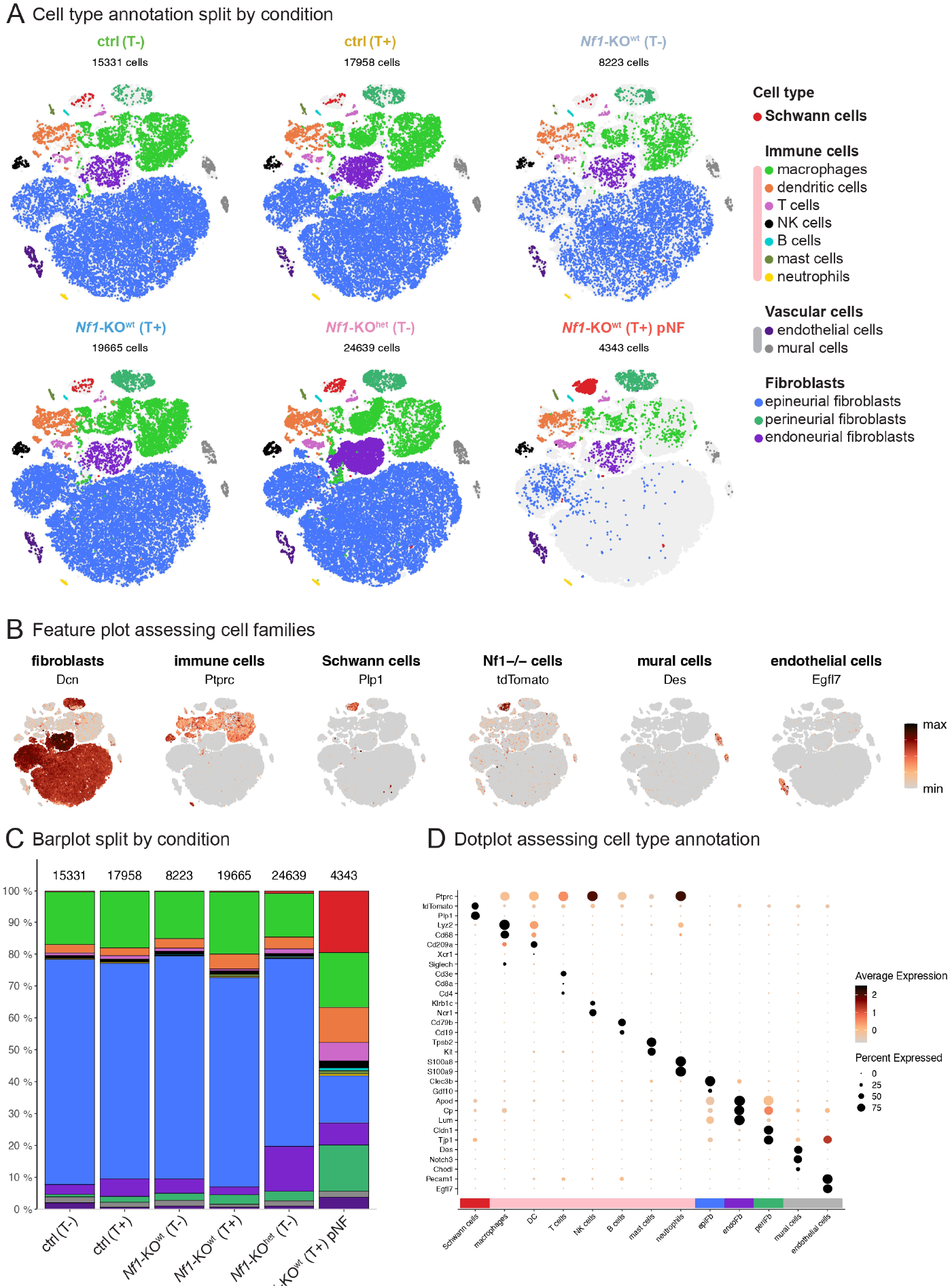
Transcriptomic atlas of control and *Nf1*-KO subcutaneous nerves. **A**. tSNE visualization of the main dataset split by experimental condition and colored according to cell types. A light grey background corresponds to cells from other conditions. Except for *Nf1*-KO^wt^ T-(n = 1), each experimental condition consists of two independent replicates. The number of cells in each condition is indicated. **B**. tSNE representation of gene expression levels to assess cell type assignment. Expression levels are log-normalized and scaled within the same range for all genes. **C**. Barplot showing cell type representativeness in each experimental condition. The number of cells in each condition is indicated. **D**. Dotplot assessing cell population annotation. Dot size represents the proportion of cells expressing the corresponding marker. The dot color represents average gene expression levels for cells concerned by the dot.

**Supplementary Figure 3.**
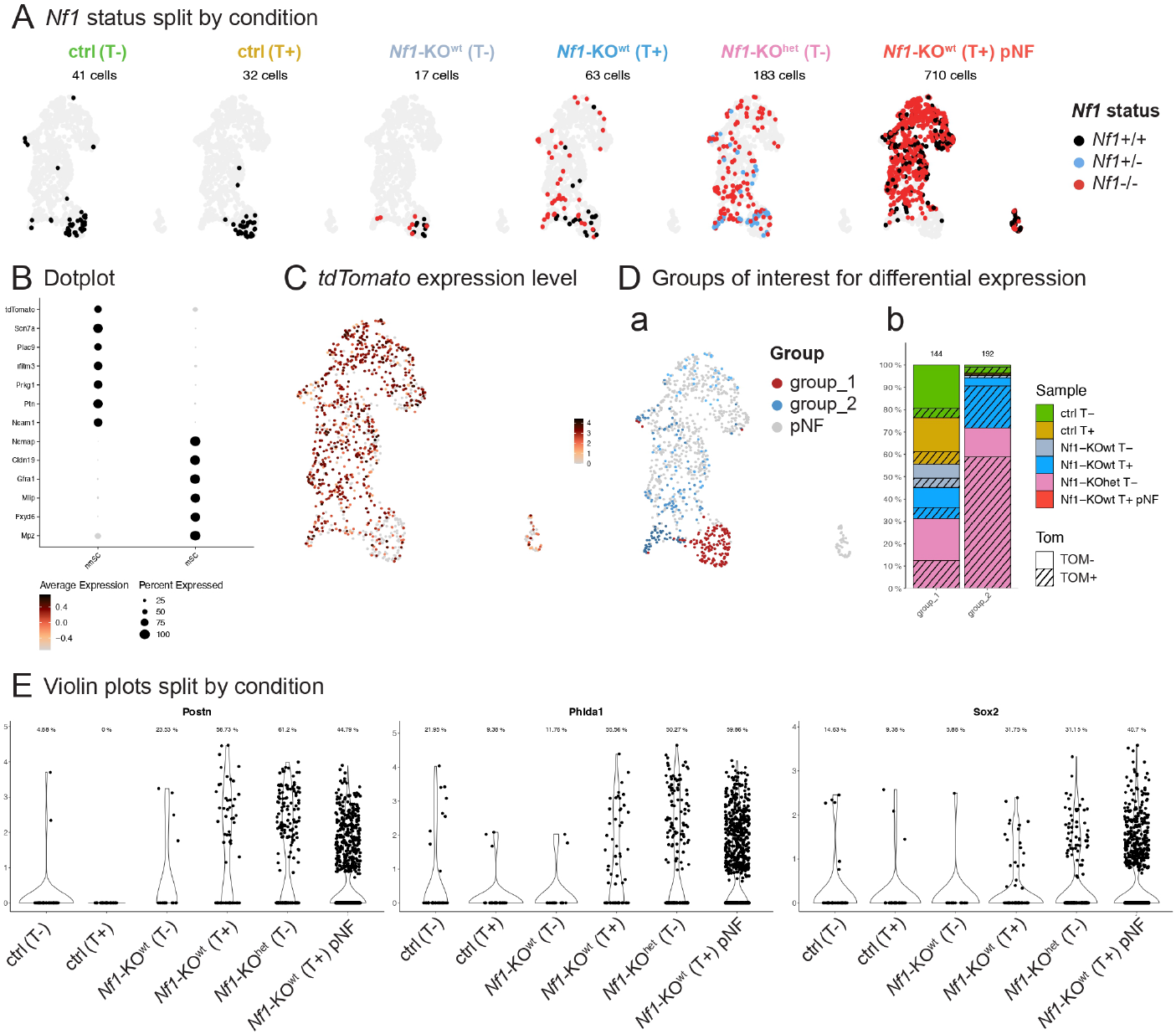
Transcriptomic signature of control and mutant Schwann cells from subcutaneous nerves. **A**. UMAP visualization of SCs dataset split by experimental condition and colored according to *Nf1* status. A light grey background corresponds to cells from other conditions. Except for *Nf1*-KO^wt^ T-(n = 1), each experimental condition consists of two independent replicates. The number of cells in each condition is indicated. **B**. Dotplot assessing cell population annotation. Dot size represents the proportion of cells expressing the corresponding marker. The dot color represents average gene expression levels for cells concerned by the dot. **C**. UMAP representation of Tomato expression level. expression level is log normalized. Parameter order = TRUE was set to represent cells with strong expression in the foreground. **D**. Schwann cells clustering. **(a)** UMAP visualization of SCs clustering. SCs were grouped into 3 groups. Group 1 (red) contains mainly *Tom*-cells while group 2 (blue shades) contains *Tom*+ cells, both isolated from nerves. Group 3 contains TOM+ SCs isolated from pNFs and was not used for differential expression analysis. **(b)** Barplot showing the distribution of each experimental condition across groups 1 and 2. Among each experimental condition, *Tom*+ SCs are depicted as hatched colors. Note that some *Nf1*-/-*Tom*+ cells have similar transcriptomic profiles as control (*Nf1*+/+) SCs. The number of cells in each group is indicated. **E**. Violin plot representing log-normalized expression for *Postn, Phlda1*, and *Sox2*, split by experimental condition. The proportion of SCs with not null expression levels is indicated.

